# Measurements of the Self-Assembly Kinetics of Individual Viral Capsids Around Their RNA Genome

**DOI:** 10.1101/265330

**Authors:** Rees F. Garmann, Aaron M. Goldfain, Vinothan N. Manoharan

## Abstract

The formation of a viral capsid-the highly—ordered protein shell that surrounds the genome of a virus—is the canonical example of self-assembly^1^. The capsids of many positive-sense RNA viruses spontaneously assemble from in vitro mixtures of the coat protein and RNA^2^. The high yield of proper capsids that assemble is remarkable, given their structural complexity: 180 identical proteins must arrange into three distinct local configurations to form an icosahedral capsid with a triangulation number of 3 (*T* = 3)^1^. Despite a wealth of data from structural studies^3–5^ and simulations^6–10^, even the most fundamental questions about how these structures assemble remain unresolved. Experiments have not determined whether the assembly pathway involves aggregation or nucleation, or how the RNA controls the process. Here we use interferometric scattering microscopy^11,12^ to directly observe the in vitro assembly kinetics of individual, unlabeled capsids of bacteriophage MS2. By measuring how many coat proteins bind to each of many individual MS2 RNA strands on time scales from 1 ms to 900 s, we find that the start of assembly is broadly distributed in time and is followed by a rapid increase in the number of bound proteins. These measurements provide strong evidence for a nucleation-and-growth pathway. We also find that malformed structures assemble when multiple nuclei appear on the same RNA before the first nucleus has finished growing. Our measurements reveal the complex assembly pathways for viral capsids around RNA in quantitative detail, including the nucleation threshold, nucleation time, growth time, and constraints on the critical nucleus size. These results may inform strategies for engineering synthetic capsids^13^ or for derailing the assembly of pathogenic viruses^14^.

---

We work with MS2 (Fig. 1a) because it is a non-trivial model system for understanding capsid assembly: Complete 28-nm capsids can be assembled in vitro from the coat proteins and RNA^16^; the assembled capsids have *T* = 3 (90 coat-protein dimers)^17^, such that they must compete with many possible malformed structures; and the RNA is suspected to play an important role in the assembly process^18–22^. Experiments probing assembly in bulk solution have shown that specific RNA sequences might initiate assembly by binding the first few coat proteins^22^. But because such measurements probe an ensemble of particles in possibly different stages of assembly, they can obscure important features of the assembly pathway. The equally important question of how that pathway can be derailed-leading to the often-overlooked minority of malformed structures observed in bulk assembly of bacteriophages^18^ and other viruses^23–25^-also remains unresolved.

**FIGURE 1.**
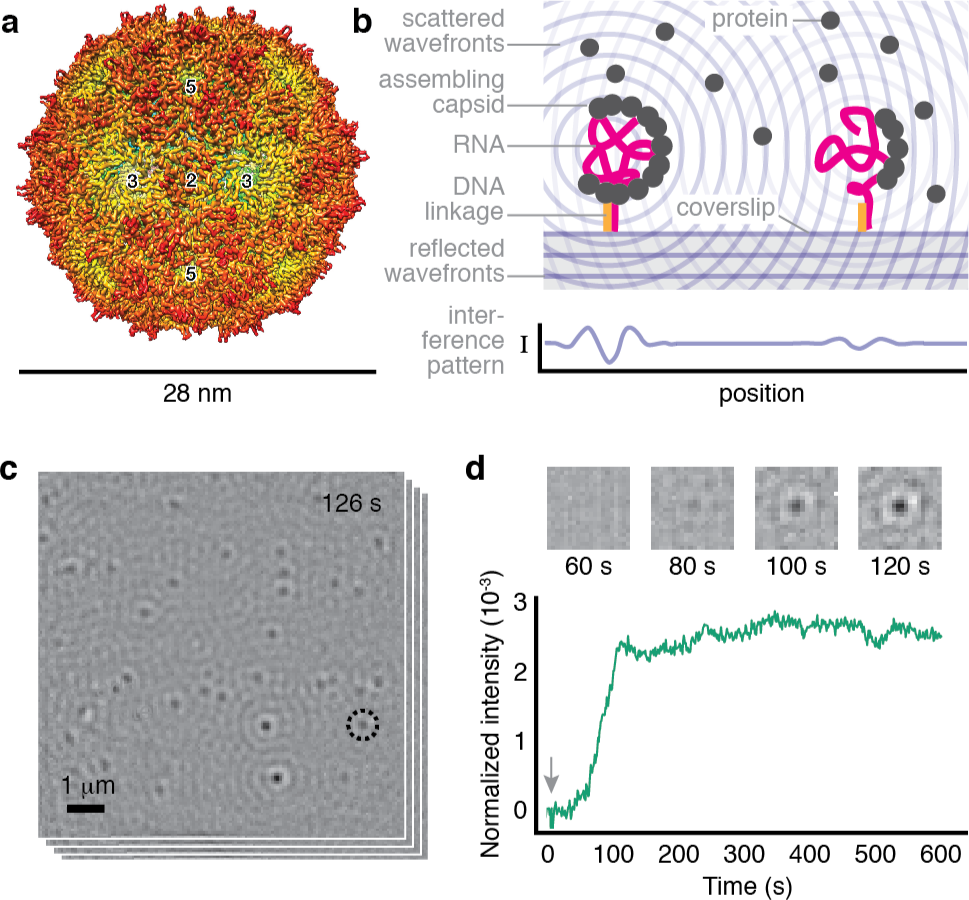
Overview of the system and measurement. (a) A structural model of the MS2 capsid from cryo-electron microscopy data (reproduced from Ref. 5) reveals its icosahedral structure with 2-, 3-, and 5-fold symmetry axes. (b) We inject a solution of MS2 coat-protein dimers over a coverslip on which MS2 RNA strands are tethered by DNA linkages^15^. As the proteins bind to the RNA, the resulting particles scatter light. Owing to destructive interference between the scattered light and a reference beam, the particles appear as dark, diffraction-limited spots. (c) We monitor many individual assembling particles in parallel. A typical image, taken 126 s after adding 2 µM dimers and representing an average of 1,000 frames taken at 1,000 frames/s, shows multiple spots. (d) The intensity of a spot as a function of time reveals the assembly kinetics of an individual particle. Top: time-series of images for the circled spot in (c). Bottom: kinetic trace for the same spot using a 1,000-frame average of data taken at 1,000 frames/s. The arrow indicates when we inject the coat protein.

Our interferometric scattering experiments address these questions because they probe the assembly of individual capsids (Fig. 1b-d). As described in Methods, each assembling particle produces a diffraction-limited spot in the field of view. Because continuous background correction in our measurement renders the RNA invisible, the final signal depends only on the number of proteins in the assembling particle. Thus, the time trace of the intensity for each spot gives a direct measure of the assembly kinetics of an individual particle.

We must measure the intensity of each spot with both high sensitivity and high dynamic range, because the capsids scatter weakly, and estimates of the assembly times range from seconds^20^ to hours^20,22^. Our apparatus (Extended Data Fig. 1) addresses both of these challenges. Because the scattering is elastic, we can use high illumination intensities with minimal risk of photodamage, enabling temporal resolutions of 1 ms. To simultaneously achieve durations of 900 s, we actively stabilize the microscope in all three dimensions, ensuring that the signal from the assembling capsid is larger than the noise due to drift. The sensitivity is then limited by shot noise. With a 1-s moving average, as shown in Fig. 1d, the peak-to-peak fluctuations from shot noise correspond to the intensity of six coat-protein dimers.

In a single experiment, we measure kinetic traces for many assembling particles in parallel, and we characterize the shape of each trace as well as variations among traces. When we inject 2 µM coat-protein dimers, we find that most traces have an initial plateau at a low intensity followed by a rapid rise and a second plateau at higher intensity (Fig. 2a and Extended Data Fig. 2). A few traces show intermediate plateaus. Most (40 out of 56) plateau at an intensity consistent with that of a full, wild-type capsid (Extended Data Fig. 3), 7 at a slightly lower intensity, and 9 at a significantly higher intensity. No such traces are observed when RNA is not tethered to the coverslip (Supplementary Information). Furthermore, negatively stained transmission electron microscopy (TEM) images reveal that most of the structures assembled in control experiments are proper capsids, with a few partial capsids and larger structures visible (Fig. 2b and Extended Data Fig. 4). We therefore infer that capsids can indeed assemble around tethered RNA strands, and that traces that reach intensities similar to those of wild-type capsids represent the formation of complete or nearly-complete capsids.

**FIGURE 2.**
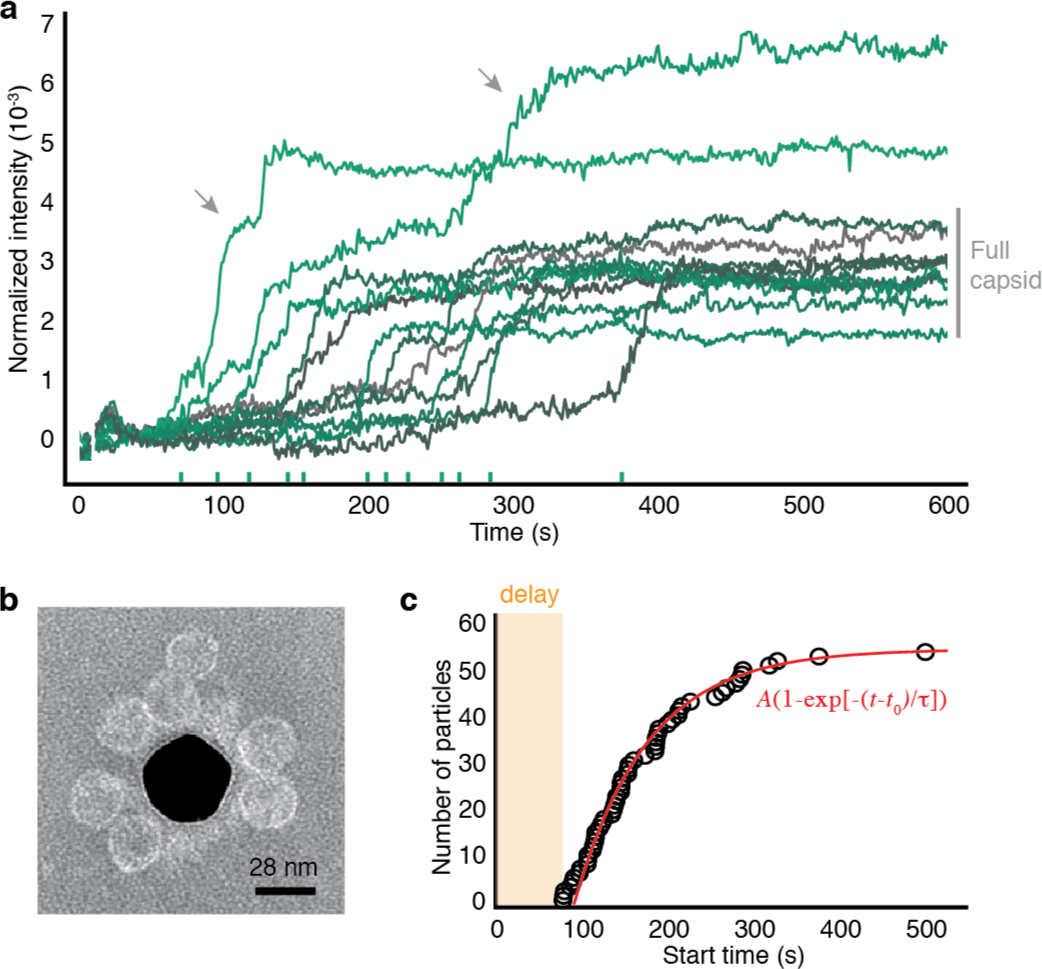
Assembly of 2 µM coat-protein dimers around surface-tethered RNA strands. (a) Kinetic traces for 12 randomly chosen particles. Ticks on the x-axis show the start times. Grey bar indicates the intensity range corresponding to wild-type capsids. Arrows point to two traces corresponding to overgrown particles. (b) A negatively stained TEM image of particles assembled around RNA strands tethered to a gold nanoparticle (dark region at center). We use a nanoparticle as the substrate because TEM cannot image through a coverslip. (c) The cumulative distribution of start times in the traces is well fit by an exponential function with *A* = 56.62 ± 0.02, *t*_0_ = 91.8 ± 0.2 s, and *τ* = 84.3 ± 0.2 s. Uncertainties in the time measurements are smaller than the diameter of the circles.

**FIGURE 4.**
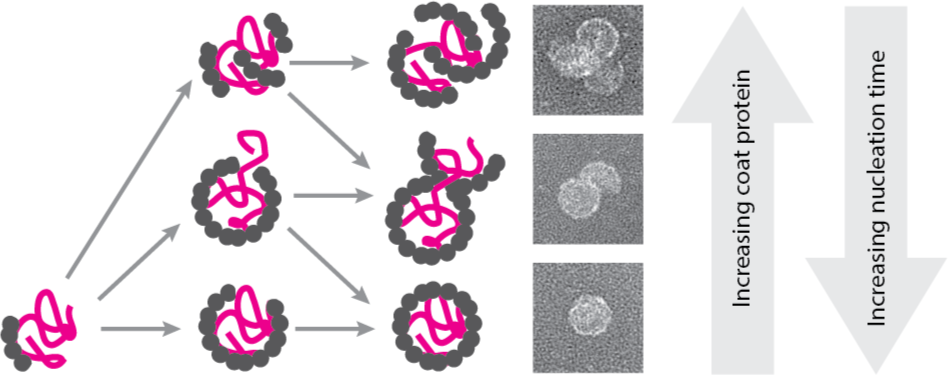
Cartoon of the inferred assembly pathways. First, a nucleus of coat proteins forms on the RNA. Bottom row: At low protein concentration, no additional nuclei form, and the nucleus grows into a proper capsid. Middle row: At higher concentrations, a second nucleus can form on an unpackaged part of the RNA, leading to a multiplet structure consisting of a nearly-complete capsid attached to a second partial or full capsid. Top row: At even higher concentrations, multiple nuclei can form and grow, leading to a monster structure consisting of many partial capsids. Example TEM images of the endpoints of each pathway are shown at right.

With this understanding, we examine what the traces reveal about the assembly pathway. A key observation is that assembly is not synchronous: the ‘start time’, the time at which the intensity rapidly increases, varies from particle to particle (Fig. 2a). We find that the cumulative distribution of start times *t* is fit well by an exponential function *A* (1 − exp [−(*t − t*_0_)*/τ*]) (Fig. 2c), where *A* is the plateau value, *t*_0_ is the delay before the start time of the first particle, and *τ* is the characteristic time (see Methods and Extended Data Fig. 5.)

The delay likely results from the combination of diffusion and a concentration threshold for assembly. We know such a threshold exists because we see no assembly when we inject 1 µM coat-protein dimers (Extended Data Fig. 6). Because the threshold is between 1 and 2 µM coat-protein dimers, we expect the delay to be of the order of the characteristic time for protein to diffuse from the 2 µM injected fluid to the surface. Indeed, that time scale is 30-55 s (Supplementary Information), and the observed delay time is 92 s.

The distribution of start times, however, does not appear to result from diffusion. The distribution is broad, with the largest start time (500 s) an order of magnitude larger than the delay time. The distribution could result from diffusion-limited growth only if the protein concentration around each RNA were to vary across the 10-µm field of view. But the time for a dimer to diffuse 10 µm is only 1 s, much shorter than the median start time. Furthermore, we estimate that about 1,000 coat-protein dimers are within 1 µm of each RNA after the initial delay. At this concentration, the pool of coat proteins is not significantly depleted by assembly, and fluctuations in concentration are negligible. We conclude that the observed kinetic traces do not result from variations in protein concentration.

Taken together, these findings rule out a diffusion-limited aggregation pathway and point strongly to nucleation and growth. The exponential shape of the cumulative distribution of start times suggests a well-defined free-energy barrier to nucleation with a nucleation time of *τ*. Although nucleation models have been used to describe the bulk assembly kinetics of empty capsids^26–28^, and computer simulations have explored nucleated pathways for capsid assembly around RNA^8–10^, direct experimental evidence for nucleation has remained elusive. The evidence that we present—the distribution of start times—cannot easily be extracted from bulk experiments^22^, which average over an ensemble of particles, or from structural experiments^29^, which have coarse temporal resolution.

Fluctuations in the intensity reveal further information about the nucleation event. Before the start time, the fluctuations are consistent with those expected from shot noise, which, as noted above, corresponds to six dimers at 1-s averaging. This measurement indirectly constrains the critical nucleus size: we can infer that sub-critical nuclei smaller than six dimers do not survive for longer than 1 s.

Additional nucleation events may be responsible for assembly going awry in some of the capsids. Most of the traces with final plateau intensities higher than that of a full capsid also show intermediate plateaus at intensities consistent with that of a full capsid (Fig. 2a). Such traces suggest that the particle undergoes a second nucleation event after the first capsid is nearly complete.

To test this hypothesis, we measure the kinetics at different concentrations of protein (Fig. 3a, and Extended Data Figs. 7 and 8). We find that the nucleation time decreases with increasing protein concentration, from about 160 s at 1.5 µM dimers to about 11 s at 4 µM (Fig. 3b). This decrease is accompanied by an increase in the fraction of overgrown particles, from 5% at 1.5 µM dimers to over 40% at 4 µM (Fig. 3c). TEM images of assembly reactions around untethered RNA (Extended Data Fig. 9) show overgrown particles with sizes corresponding to the final intensities seen in the kinetic traces. Many of the overgrown particles consist of bunches of partial or nearly-complete capsids (Fig. 3d).

**FIGURE 3.**
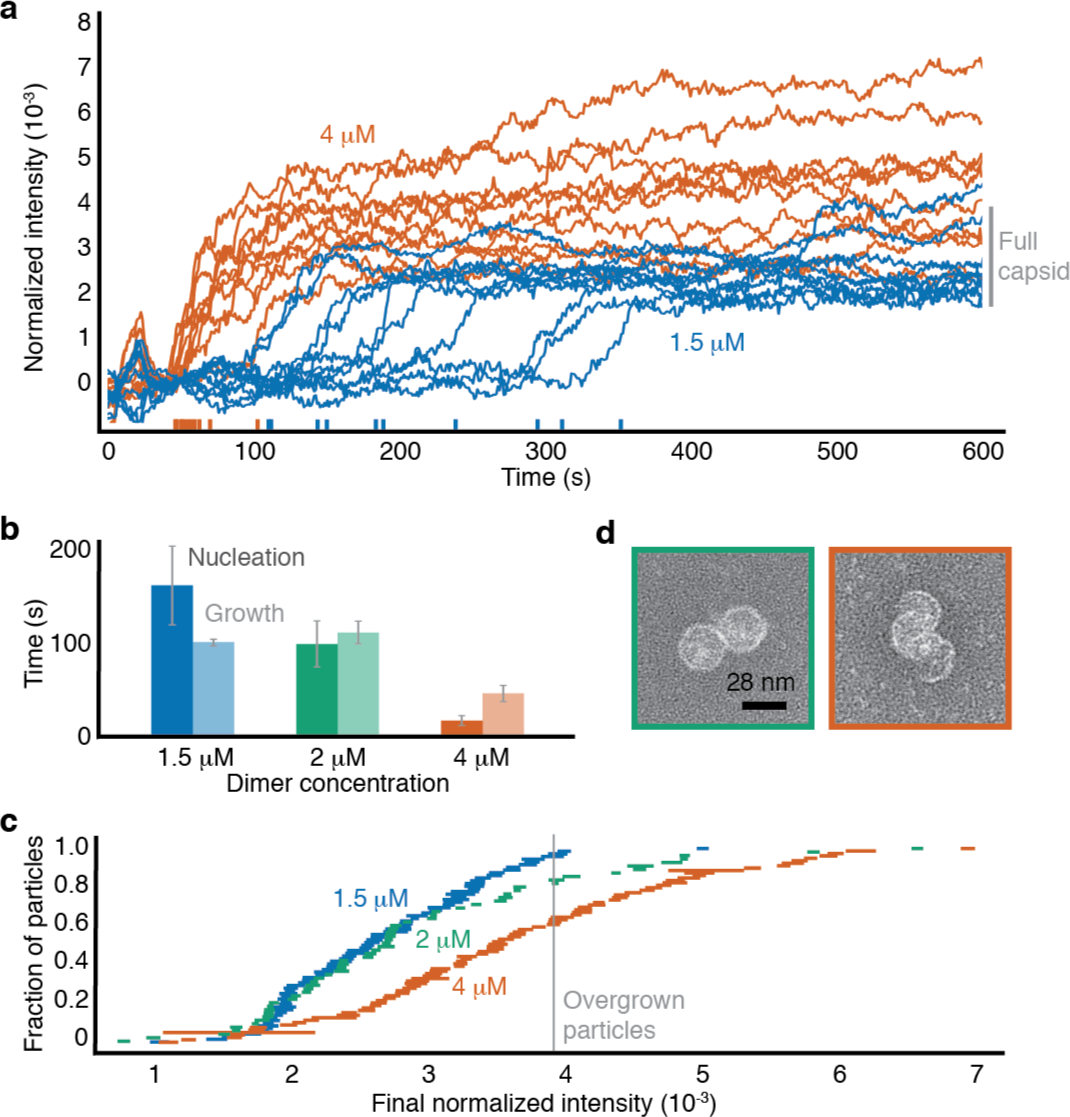
Assembly kinetics at different protein concentrations. (a) Kinetic traces for 10 randomly chosen particles at 1.5 µM and 4 µM coat-protein dimers. (b) Measured nucleation times and median growth times. Error bars represent the standard deviation from three experiments. (c) Cumulative distributions of the final intensities show that the fraction of overgrown particles increases with protein concentration. The length of each horizontal bar is the standard deviation calculated from the last 50 s of each trace. (d) TEM images of overgrown particles around untethered RNA. Left: an attached pair of nearly complete capsids at 2 µM protein. Right: connected partial capsids at 4 µM protein.

The kinetic traces and images of the overgrown structures suggest a pathway involving more than one nucleation event. However, many of the traces at 4 µM coat-protein dimers do not show intermediate plateaus (Fig. 3a). To understand why, we measure how long it takes a particle to reach the intensity of a full capsid after it starts growing (see Methods). We find that these ‘growth times’ decrease with increasing protein concentration, but less rapidly than do the nucleation times (Fig. 3b). When the nucleation time is smaller than the growth time, as it is in experiments with 4 µM dimers, additional nuclei can form before the first has time to grow. Under such conditions, most of the kinetic traces should not—and indeed, do not—show intermediate plateaus.

Thus, the viral RNA creates a competition between nucleation and growth, as sketched in Fig. 4. Similar scenarios have been observed in computer simulations of capsid assembly on polymer scaffolds^8,9^, and may explain the formation of the ‘monster’^23^ and ‘multiplet’^24,25^ structures observed in experiments with other viruses. These structures are not observed in experiments on the assembly of empty capsids^30,31^, confirming that the RNA plays a critical role in the assembly pathways.

Our individual-particle measurements also rule out some competing pathways. The assembly of proper capsids appears to follow a one-step nucleation pathway rather than a multi-step one^32^. Also, the formation of overgrown structures appears to result from multiple nucleation events rather than the ‘spiraling’ pathway^6,7^ observed in local rules-based simulations, or the ‘en-masse’ pathway^8,9^ observed in Brownian dynamics simulations. Because our measurements involve thermally annealed RNA (see Methods), they do not yet resolve whether the pathway is fine-tuned by local folding patterns in the viral RNA^33,34^.

While our observations are specific to in vitro assembly, the observed threshold concentration for nucleation supplies a hypothesis for viral replication in vivo. Because viral RNA that is inside a capsid cannot be replicated or translated, a virus such as MS2 must delay encapsidation until its components have been produced in sufficient quantities. With a threshold for nucleation, assembly (and encapsidation) would take place only after there is enough viral RNA and coat protein to form many new virus particles.

Although we expect the assembly pathways to differ for different viruses and buffer conditions, our measurements of the nucleation time, threshold, growth time, and subcritical fluctuations in MS2 provide important constraints on models of assembly. As a result, the structures of the assembly intermediates and the critical nucleus, which have long eluded direct imaging methods, might now be inferred through quantitative comparisons of simulated^8,9^ and measured individual-particle kinetics. This approach might identify conditions for assembly of synthetic viruses^13^ and new targets for antiviral therapies that work by disrupting capsid assembly^14^.

## METHODS

### Interferometric scattering microscope

Our microscope is configured in wide-field mode and is similar to the setup described by Ortega-Arroyo and coworkers^35^. A 450 nm, 100 mW, single-mode diode laser (PD-01251, Lasertack) illuminates the sample. The current driving the laser is modulated with a square wave at a frequency of 1 MHz to decrease the coherence of the laser and limit intensity variations in the background^36^. The beam (shown in blue in Extended Data Fig. 1) is spatially filtered by a polarization-maintaining single-mode optical fiber (fiber 1; PM-S405-XP, Thorlabs). The filtered light is collected by a lens (lens 1; achromatic doublet, focal length = 25 mm, Thorlabs), reflected from a polarizing beamsplitter cube (CCM1-PBS251, Thorlabs), and focused onto the back aperture of the objective (100× oil-immersion, 1.45 NA Plan Apo *λ*, Nikon) to produce collimated illumination in the imaging chamber. The light that is backscattered from the sample and light that is reflected from the water-coverslip interface are collected by the objective and imaged onto camera 1 (MV1-D1024E-160-CL, Photon Focus) by the tube lens (achromatic doublet, focal length = 300 mm, Thorlabs). We use achromatic half and quarter-wave plates (AHWP3 and AQWP3, Bolder Vision Optik) with the polarizing beamsplitter to make an optical isolator that minimizes the intensity lost at the beamsplitter. The total magnification is 150×, such that each pixel on the camera views a field of 70 nm. All images are recorded with a bit depth of 12.

The illumination intensity, set to approximately 3 kW/cm^2^ when we record data at 1,000 Hz and 0.3 kW/cm^2^ at 100 Hz, is similar to that typically used in single-molecule fluorescence experiments^37^. To minimize any possible radiation damage, we use an exposure time that is almost equal to the total time between frames, and we dim the imaging beam with absorptive filters so that the camera pixels are nearly saturated. The total field of view is 140 pixels × 140 pixels (9.8 µm × 9.8 µm) at 1,000 Hz and 200 pixels × 200 pixels (14 µm × 14 µm) at 100 Hz.

We use short-wavelength light (*λ* = 450 nm) because the intensity of the image scales with *λ^−^*^2^. While shorter wavelength lasers are available, we find that they can damage both the sample and optical components when used at high intensities. Control experiments at different illumination intensities are described below and in Supplementary Information, which also contains additional notes on the configuration of the microscope.

The intensity of each diffraction-limited spot is approximately linearly proportional to the number of proteins bound to the RNA strand. The intensity of a spot is 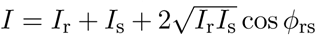, where *I*_r_ is the intensity of the reflected wave, *I*_s_ the intensity of the scattered wave, and *φ*_rs_ the phase difference between the two. The term *I*_s_ can be neglected since the scattered light is dim compared to the reflected light, so the normalized intensity *I*_norm_ = *I/I*_r_ − 1 is proportional to the total polarizability of the assembling particle^12^, which is approximately the sum of a protein component and an RNA component. Because the RNA component is static, it is part of the background, which is subtracted. As a result, the normalized intensity is linearly proportional to the number of proteins in the assembling particle.

### Active stabilization

The position of the coverslip relative to the objective is actively stabilized to a few nanometers in all three dimensions. Each dimension is controlled separately through a proportional control loop on the PC. During each iteration of the loop, the position of the coverslip is measured, and the voltage driving the piezoelectric actuators is modified to keep the coverslip in its original position.

The height of the coverslip above the objective is measured by tracking the position of a laser (red in Extended Data Fig. 1) that is totally internally reflected by the coverslip-water interface, as described by Ortega-Arroyo and coworkers^35^. We use a 785-nm, 90 mW, single-mode diode laser (L785P090, Thorlabs) that is coupled through a single-mode fiber (fiber 2; S630-HP, Thorlabs). The laser is driven with a constant current (27 mA) that is well below threshold (35 mA), which we find improves the intensity stability of the laser. After exiting the optical fiber, the beam is collected by lens 2 (plano-convex, focal length = 20 mm, Thorlabs), reflects from a dichroic mirror (700-nm short-pass, Edmund Optics), and is focused onto the back aperture of the objective. We align the beam so that after exiting the objective, it totally internally reflects from the coverslip-water interface and re-enters the objective. The total power incident on the coverslip is less than 1 µW. The return beam reflects from the coverslip and then from a D-shaped mirror (Thorlabs) and is detected with camera 2 (DCC1545M, Thorlabs). A long-pass filter (700-nm, Thorlabs, not shown in Extended Data Fig. 1) attenuates any light from the imaging beam that is also incident on camera 2. When the height of the coverslip changes, the return beam is displaced laterally across camera 2, resulting in a change in the measured center-of-brightness. Under active stabilization, any changes in the center-of-brightness are measured and corrected every 30 ms.

The in-plane position of the coverslip is measured by tracking a 30-nm gold particle that is adsorbed to the coverslip surface (see next subsection for details of how we prepare the coverslips). Before each experiment, we find one of the adsorbed gold particles by looking for spots that have a normalized intensity of approximately 0.2. We then move the coverslip so that the spot is near the edge of the field of view. Using a 16 × 16-pixel region of the field of view, we record a static background image of the coverslip with no particles present and then move the gold particle into the center of this small field of view. Before tracking the position of the gold particle, we process its image in the small field of view by subtracting off the static background, applying a bandpass filter (passing features of size 1 to 7 pixels) to smooth the image, and taking the time-median of 33 images of the particle (recorded at 33 Hz) to reduce shot noise. We then use the program Trackpy^38^ to locate the position of the particle. We use this position for the active stabilization loop, which runs once per second. The in-plane control loop frequency (1 Hz) is lower than that of the out-of-plane control loop (33 Hz) because of the time required to collect the median image of the particle.

The active stabilization loops are implemented in a Python script (http://github.com/manoharan-lab/camera-controller). The same script includes a real-time image processing routine that allows us to see growing MS2 particles while collecting data.

### Coverslip and gold nanoparticle functionalization

We adapt the protocols described by Joo and Ha^39^ to coat glass coverslips with a layer of PEG molecules, about 1% of which are functionalized with short DNA oligonucleotides. We find that many brands of #2 coverslips are unsuitable for assembly measurements because they have imperfections that scatter too much light. We use only #2 thickness, 24 mm × 60 mm rectangular glass microscope coverslips from Globe Scientific, Inc. Details of how we functionalize the coverslips and decorate them with 30-nm gold particles are in Supplementary Information.

### Flow cell design and construction

We build chips that each contain 10 separate flow cells above a single coverslip. Each chip consists of two sheets of cut, clear acrylic that are sealed together and to the coverslip with melted Parafilm (Bemis). Each flow cell has an imaging chamber that is used for the assembly experiments, an inlet cup to hold fluid before it is introduced into the imaging chamber of the flow cell, a short inlet chamber to connect the inlet cup to the imaging chamber, and an outlet chamber. We use acrylic, a hard plastic, because we find that soft materials such as polydimethylsiloxane lead to more warping of the coverslip during injection of the protein. A detailed description of the flow cells and their construction is given in Supplementary Information.

### Growth of MS2 and purification of its coat protein and RNA

We grow wild-type MS2 by infecting liquid cultures of E. coli strain C3000 (a gift from Peter Stockley at the University of Leeds) and purifying the progeny viruses following the protocols of Strauss and Sinsheimer^40^. We purify coat protein from the virus particles following the cold acetic acid method described by Sugiyama, Hebert, and Hartmann^16^. We purify RNA from freshly grown MS2 virions using an RNA extraction kit (RNeasy, Qiagen). Details about how we assess the purity of these materials are described in Supplementary Information.

We store the purified virus particles at 4 °C and discard them after about 1 month. We store the protein at 4 °C and discard it after 1 week. We store the RNA at *−*80 °C and discard it after about 1 year.

### Surface-immobilization of MS2 RNA by DNA linkages

To immobilize MS2 RNA at the coverslip surface, we first hybridize the 5’-end of the RNA to a 60-base-long linker oligo (Integrated DNA Technologies). The 40 bases at the 5’-end of the linker are complementary to the 40 bases at the 5’-end of the RNA, and the remaining 20 bases are complementary to the sequence of the surface oligo (Extended Data Fig. 1). To anneal the linker to the MS2 RNA, we add a 10-fold molar excess of the linker oligo to 500 nM MS2 RNA in hybridization buffer (50 mM Tris-HCl, pH 7.0; 200 mM NaCl, 1 mM EDTA), heat the mixture to 90 °C for 1 s, and then cool it to 4 °C at a rate of −1 °C/s. Excess linker is removed with a 100-kDa-MWCO centrifugal filter unit (EMD Millipore) at 14,000 g. The 60-base-long oligonucleotides do not pass through the filter; instead, they stick to the membrane. We confirm RNA-DNA binding by native 1% agarose gel electrophoresis (Supplementary Information). We confirm that the RNA-DNA constructs specifically bind our DNA-functionalized coverslips by interferometric scattering microscopy (Supplementary Information). The sequence of the linker is 5’-CGACAGGAAGTTGAGCAGGACCCCGAAAGGGGTCCCACCCAACCAACCAACCAACCAACC-3’.

### Calibration experiment

We measure the intensities of MS2 RNA and wild-type MS2 virus particles (Extended Data Fig. 3) by imaging the particles as they adsorb to an APTES-functionalized coverslip. For these experiments we do not use a flow cell. Instead, we use a ‘lean-to’ sample chamber^41^ made of 1-mm-thick glass slides (Micro Slides, Corning) that are cut, cleaned by pyrolysis (PYRO-CLEAN, Tempyrox Co.), and sealed in place with vacuum grease (High vacuum grease, Dow Corning). To perform the calibration experiment, we first fill the sample chamber with TNE buffer (50 mM Tris-HCl, pH 7.5; 100 mM NaCl; 1 mM EDTA) and focus the microscope onto the coverslip. We then exchange the buffer in the sample chamber with a solution containing both MS2 RNA and wild-type MS2 virus particles at a concentration of 0.1 nM each in TNE buffer. We record movies (100 Hz) of these particles nonspecifically adsorbing to the coverslip.

We see two well-separated populations in the distribution of intensities of the particles that bind (Extended Data Fig. 3). We assume that the lower-intensity population is due to the RNA strands and the higher-intensity population is due to the MS2 viruses. To determine the median and width of each intensity population, we separate the two using an intensity threshold (0.003) that lies between them.

### Assembly experiments

For assembly experiments, we fill a flow cell with hybridization buffer containing 0.2% Tween-20 (Sigma-Aldrich) and let it sit for 10 min. We find that this 10-min incubation with Tween-20 prevents the MS2 coat protein from adsorbing to the coverslip through defects in the PEG layer. Next, we flush out the Tween-20 with fresh hybridization buffer, find the center of the imaging chamber, focus the microscope onto the coverslip, and begin the out-of-plane active stabilization control loop. Then we locate a 30-nm gold particle within 50 µm of the center of the imaging chamber and start the in-plane active stabilization control loop. With the setup actively stabilized in all three dimensions, we inject 1 nM RNA-DNA complexes in hybridization buffer and record a short movie of them adsorbing to the coverslip. After 10–100 complexes bind, we flush the imaging chamber by pumping 120 µL of assembly buffer through the chamber over the course of 12 min. Then we start recording a movie and inject the coat-protein dimers in assembly buffer. The injection starts 4 s into the movie.

### Image processing

We process the images to normalize them and to reduce fluctuations in the background intensity. We apply an approach similar to the ‘pseudo-flat-fielding’ method described by Ortega-Arroyo and coworkers^35^. The images in Fig. 1, and Extended Data Fig. 3, are processed in this way, as are all the movies included in the Supplementary Information.

Each raw image, denoted *I*_raw_, is processed according to the following steps: First, a dark image, *I*_dark_, is acquired by taking the time-median of many frames (200 frames for 100 Hz data and 2,000 for 1,000 Hz data) when the illumination beam is blocked. This image is subtracted from each raw image, yielding *I*_bkgd_ = *I*_raw_−*I*_dark_. Second, features bigger than *σ*_1_ = 1.5 pixels are removed by subtracting a Gaussian blur, yielding *I*_smooth_ = *I*_bkgd_— blur(*I*_bkgd_*, σ*_1_), where blur(*I, σ*) is 2D Gaussian blur of the image, *I*, using a standard deviation *σ*. We choose *σ*_1_ = 1.5 to minimize intensity changes that arise from time-varying background fringes, even though this choice slightly decreases the normalized intensities of the particles on the coverslip. Third, the image is normalized to the background that has been blurred with *σ*_2_ = 20 pixels, so that particles on the coverslip and stray fringes smaller than *σ*_2_ do not affect the normalization. This process yields *I*_norm_ = (*I*_smooth_)*/*blur(*I*_bkgd_*, σ*_2_). Because each image is normalized independently of other images in the time-series, fluctuations in the illumination intensity in time do not affect *I*_norm_. Finally, all remaining static features in the background are removed by subtracting the time-median of many frames (300 frames for 100 Hz data and 3,000 for 1,000 Hz data) of the movie, yielding the final processed image *I*_final_ = *I*_norm_ − *I*_norm,mod_. The noise in *I*_final_ is set by shot noise for the first few seconds after the background subtraction, but after this time, fluctuations in the background intensity due to uncorrected mechanical drift are the main source of measurement noise.

### Identifying and measuring assembling particles

To identify assembling particles, we manually locate the centers of all dark spots that appear and are between 1 and 4 pixels across in each processed interferometric scattering movie. We repeat this procedure multiple times using different frames for the background subtraction to ensure that no dark spots are missed. For each of these spots, we measure the mean intensity in a circle of radius 1 pixel that is centered on the particle as a function of time.

We reject any spot that: (1) instantaneously appears in the movie, indicating that it is from a particle that has adsorbed to the coverslip; (2) is near the gold particle used for active stabilization or near a defect on the coverslip that has comparable intensity (greater than 0.1); (3) is near a particle that adsorbs to or desorbs from the coverslip, such that its intensity is altered by the particle; (4) is so close to another spot that the interference fringes of the two spots overlap; (5) is near the edge of the field of view; or (6) grows at a slow and consistent rate over the course of the measurement, consistent with protein assembly in the absence of RNA. We describe how each of these criteria are applied in the Supplementary Information.

### Determining start and growth times

The cumulative distribution functions of the start times before assembly (Figs. 2 and Extended Data Figs. 5) are measured as follows. Each start time is defined as the time at which a kinetic trace reaches an intensity of 0.001. To measure this time, we smooth each trace using a 1,000-frame moving average. The first time that the smoothed trace reaches an intensity greater than 0.001 is called *t*_1_, and the last time that the smoothed trace has an intensity less than 0.001 is called *t*_2_ (ignoring any late detachment events or drifts in intensity). The start time is then determined as *t*_start_ = (*t*_1_ + *t*_2_)*/*2. To estimate the uncertainty in each start time, we calculate the half-width of the moving-average window and (*t*_2_ − *t*_1_)*/*2, and we take the greater of the two. The cumulative distribution function of start times is obtained by sorting the measured values of *t*_start_.

We then fit the cumulative distribution to the exponential function *N* (*t*) = *A* (1 − exp [− (*t − t*_0_)*/τ*]) using a Bayesian parameter-estimation framework. A uniform, unbounded prior is used for all parameters. The exponential function is first inverted, yielding

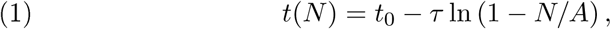

where the fit parameters are *t*_0_, *A*, and *τ*. The posterior probability distribution *p*(*t*_0_*, A, τ| D*_CDF_*, M*), where *D*_CDF_ is the observed cumulative distribution function and *M* is the model (Equation (1)), is then sampled using an affine-invariant ensemble Markov-chain Monte Carlo sampler^42^ with 50 walkers that take 500 steps each. The walkers are initially distributed in a narrow Gaussian around the peak of the posterior probability density function. The position of the peak is calculated from a least-squares fit to *t*(*N*). The walkers reach an equilibrium distribution after approximately 200 steps. Pair plots of the positions of the walkers on every step after the burn-in are shown in Extended Data Fig. 5, along with the marginal distributions for each fit parameter. The best-fit parameters reported in the text are taken as the 50^th^ percentile of the marginal distributions, and the reported uncertainties represent a credibility interval from the 16^th^ to the 84^th^ percentile.

To determine the growth time we first take the portion of each kinetic trace that lies between the start time and the time at which the intensity first reaches the 10^th^ percentile of the capsid intensity distribution (Extended Data Fig. 3), and fit this portion of the trace to a line, using a least-squares method. We then estimate the time required to grow a full capsid (bind 90 dimers) by approximating the growth rate as the slope of the linear fit.

### Control assembly experiment with lower illumination intensity

To test whether the intensity of the incident beam affects the assembly process, we perform a set of duplicate control experiments with 2 µM coat-protein dimers and a light intensity that is 10-fold smaller (approximately 0.3 kW/cm^2^). The results, shown in the Supplementary Information, indicate that the incident light does not qualitatively affect the assembly process.

### TEM of assembled particles

We use negative staining and TEM to image the protein structures that form on MS2 RNA. First, we describe assembly experiments with MS2 RNA that is tethered to the surface of 30-nm gold particles (Fig. 2e, Extended Data Fig. 4). The surfaces of the gold particles are functionalized in a way that is similar to that used for the coverslips. The protocol is identical to that used to prepare the tracer particles for active stabilization (Supplementary Information), except that we use NHS-PEG-N_3_ instead of NHS-PEG. To conjugate DNA oligonucleotides to the PEG-coated gold particles, we add 5 µM DBCO-DNA to 10 nM gold particles in PBS without Ca or Mg. The mixture is left at room temperature overnight in a tube rotator and then washed 5 times by centrifuging the mixture at 8,000 g for 5 min and resuspending in TE buffer.

To perform the assembly reaction, we add a 100-fold molar excess of RNA-DNA complexes (20 nM) to the gold particles (0.2 nM) and equilibrate the mixture in TNE buffer for 1 hr on ice. We then take 6 µL of this mixture, add 0.42 µL of 30 µM coat-protein dimers suspended in 20 mM acetic acid, and let the mixture sit for 10 min at room temperature. The mixture is then added to a plasma-etched carbon-coated TEM gird (Ted Pella), left to sit for 1 min, and then removed by blotting with filter paper. Then 6 µL of methylamine tungstate stain solution (Nanoprobes) is added and left to sit for 1 min before removal by blotting with filter paper. We visualize the samples on a Tecnai F20 (FEI) transmission electron microscope operated at 120 kV. Images are captured on a 4,096 × 4,096-pixel CCD camera (Gatan). Representative images are shown in Extended Data Fig. 4 along with images of control reactions involving bare RNA without the DNA linkage.

We also perform assembly reactions with RNA that is free in solution. This is done by mixing varying concentrations of coat protein with 10 nM of RNA in assembly buffer. After allowing the assembly reaction to proceed for a fixed amount of time, the mixture is imaged by TEM, as described above. Representative electron micrographs of particles assembled with 1.5, 2, and 4 µM coat-protein dimers are shown in Extended Data Fig. 9.

### Buffer recipes

**Assembly buffer:** 42 mM Tris-HCl, pH 7.5; 84 mM NaCl; 3 mM acetic acid; 1 mM EDTA

**Hybridization buffer:** 50 mM Tris-HCl, pH 7.0; 200 mM NaCl; 1 mM EDTA

**TAE buffer:** 40 mM Tris-acetic acid, pH 8.3; 1 mM EDTA

**TNE buffer:** 50 mM Tris-HCl, pH 7.5; 100 mM NaCl; 1 mM EDTA TE buffer: 10 mM Tris-HCl, pH 7.5; 1 mM EDTA

### Code availability

The code used to analyze the data is available from the corresponding author on reasonable request.

### Data availability

The datasets generated and analyzed during the current study are available from the corresponding author on reasonable request.

## ACKNOWLEDGEMENTS

We thank Peter Stockley and Amy Barker at the University of Leeds for sending us initial stocks of MS2 virus and C3000 cells and their growth protocols. We thank Philip Kukura, Marek Piliarik, Vahid Sandoghdar, and Michael Brenner for helpful discussions. This work is supported by the Harvard MRSEC under National Science Foundation grant no. DMR-1420570. Additionally, this material is based upon work supported in part by the National Science Foundation Graduate Research Fellowship under grant no. DGE-1144152. It was performed in part at the Center for Nanoscale Systems (CNS), a member of the National Nanotechnology Coordinated Infrastructure Network (NNCI), which is supported by the National Science Foundation under NSF award no. 1541959. CNS is part of Harvard University.

## AUTHOR CONTRIBUTIONS

VNM came up with the idea of studying virus self-assembly using interferometric scattering microscopy and developed this idea with RFG and AMG. AMG and RFG designed the experimental setup, performed the experiments, and analyzed the data. VNM supervised the project. AMG, RFG, and VNM wrote the manuscript.

## EXTENDED DATA

**EXTENDED DATA FIGURE 1.**
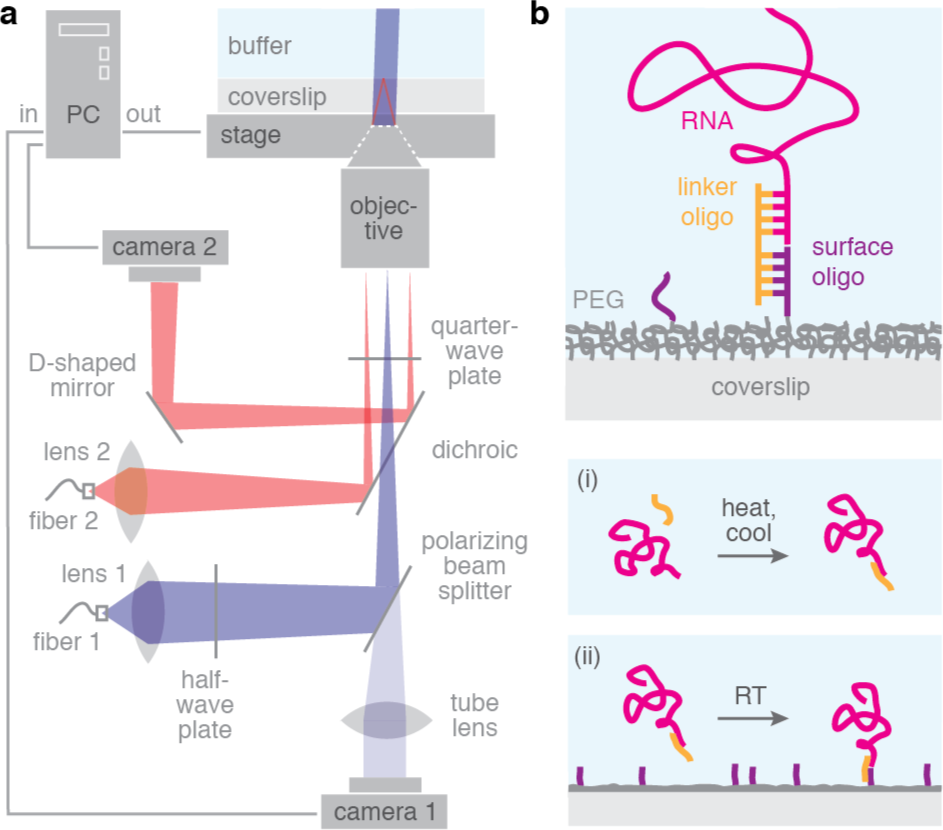
Diagram of the interferometric scattering microsco DNA linkages. (a) Our microscope is similar to the setup described by Ortega-Arroyo and coworkers^35^. 450-nm light (blue) is used for illumination. 785-nm light (red) is used for active stabilization in the dimension perpendicular to the coverslip surface. Details of the instrument are described in Methods. (b) We use DNA linkages^15^ to bind MS2 RNA to the surface of a microscope coverslip. Top: diagram of the basepairing between the 5’-end of the RNA, a linker oligo, and a surface oligo that is covalently bound to the PEG-functionalized coverslip. Bottom: to construct the linkages we (i) bind the RNA to the linker oligo in solution by thermal annealing, and then (ii) add the RNA-DNA complexes to the functionalized coverslips at room temperature. Details of the process are described in Methods.

**EXTENDED DATA FIGURE 2.**
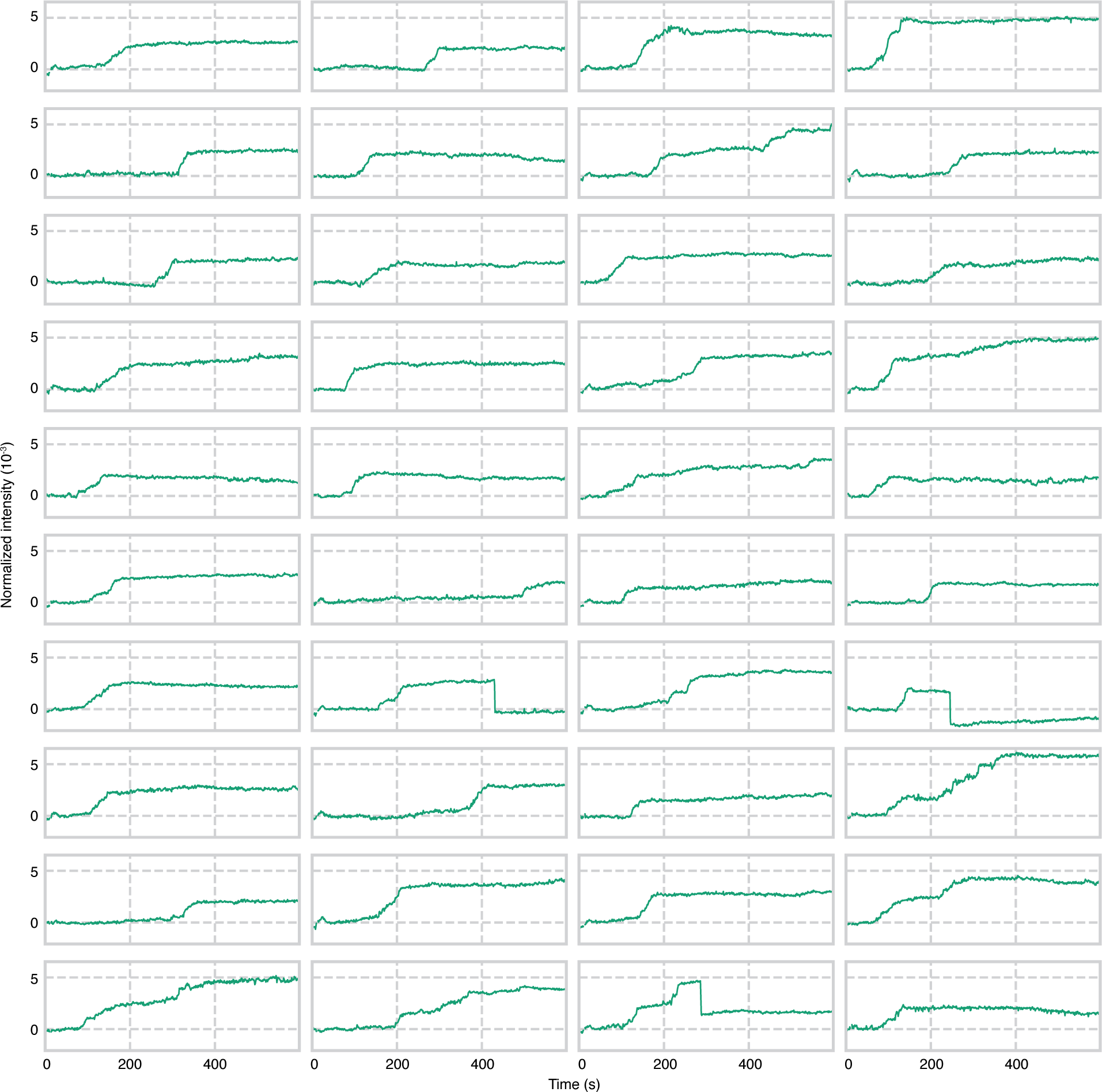
Assembly of 2 µM coat-protein dimers. Kinetic traces for 40 of the 56 observed assembling particles in one experiment are shown. Some traces show abrupt drops in intensity, which we interpret as detachment events. One of the above traces drops to an intensity of between −0.001 and −0.002, which is approximately the negative intensity of the RNA in the background image. We therefore interpret this event as the detachment of the RNA and assembled proteins from the surface. One trace drops to an intensity near 0, suggesting that the assembled protein has detached from the RNA, while the RNA remains on the surface. One of the traces drops from an intensity near 0.005 by an amount (0.0032) that corresponds to a full capsid, suggesting that overgrown particles can contain capsids. A portion of the traces from the same experiment appear in Fig. 2, and one trace from the experiment appears in Fig. 1d. The final intensities of the particles in this experiment are used for Fig. 3c. The traces are measured from the data shown in Supplementary Movie 1. The data are recorded at 1,000 Hz and are plotted with a 1,000-frame average.

**EXTENDED DATA FIGURE E3.**
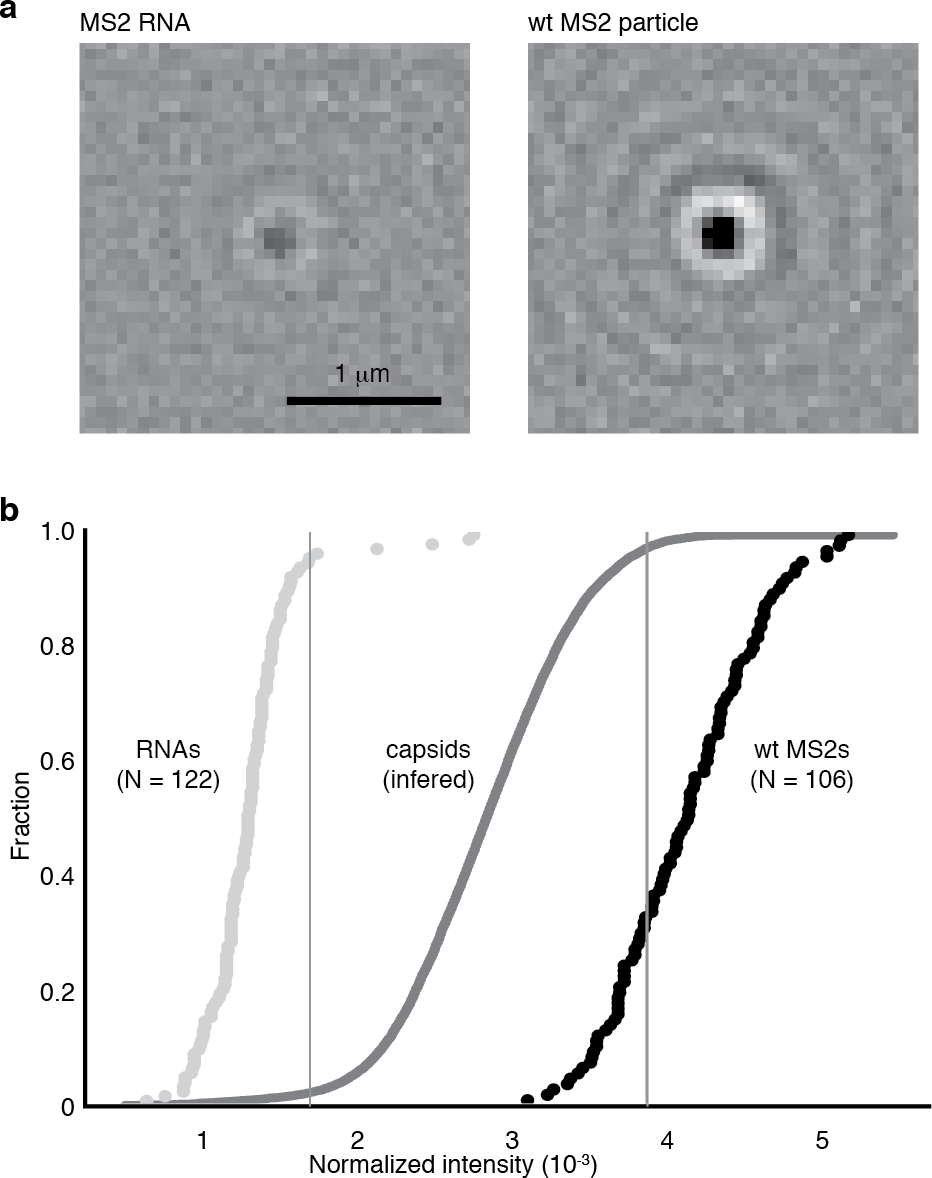
Cumulative distribution of the normalized intensities of MS2 RNA strands and wild-type MS2 virus particles measured in the interferometric scattering microscope. See Methods for details of the measurement. (a) Images of a single MS2 RNA strand (left) and a single wild-type MS2 virus particle (right). Both images are recorded at 100 Hz and shown with a 300-frame average. (b) We infer the cumulative distribution of intensities for MS2 capsids that fully assemble on surface-tethered RNA by convolving the intensity distribution of the wild-type MS2 particles with the negative of the intensity distribution of the MS2 RNA strands. The gray lines, which mark where the capsid distribution reaches 2.3% and 97.7%, denote the interval we use for identifying full capsids in the kinetic traces.

**EXTENDED DATA FIGURE E4.**
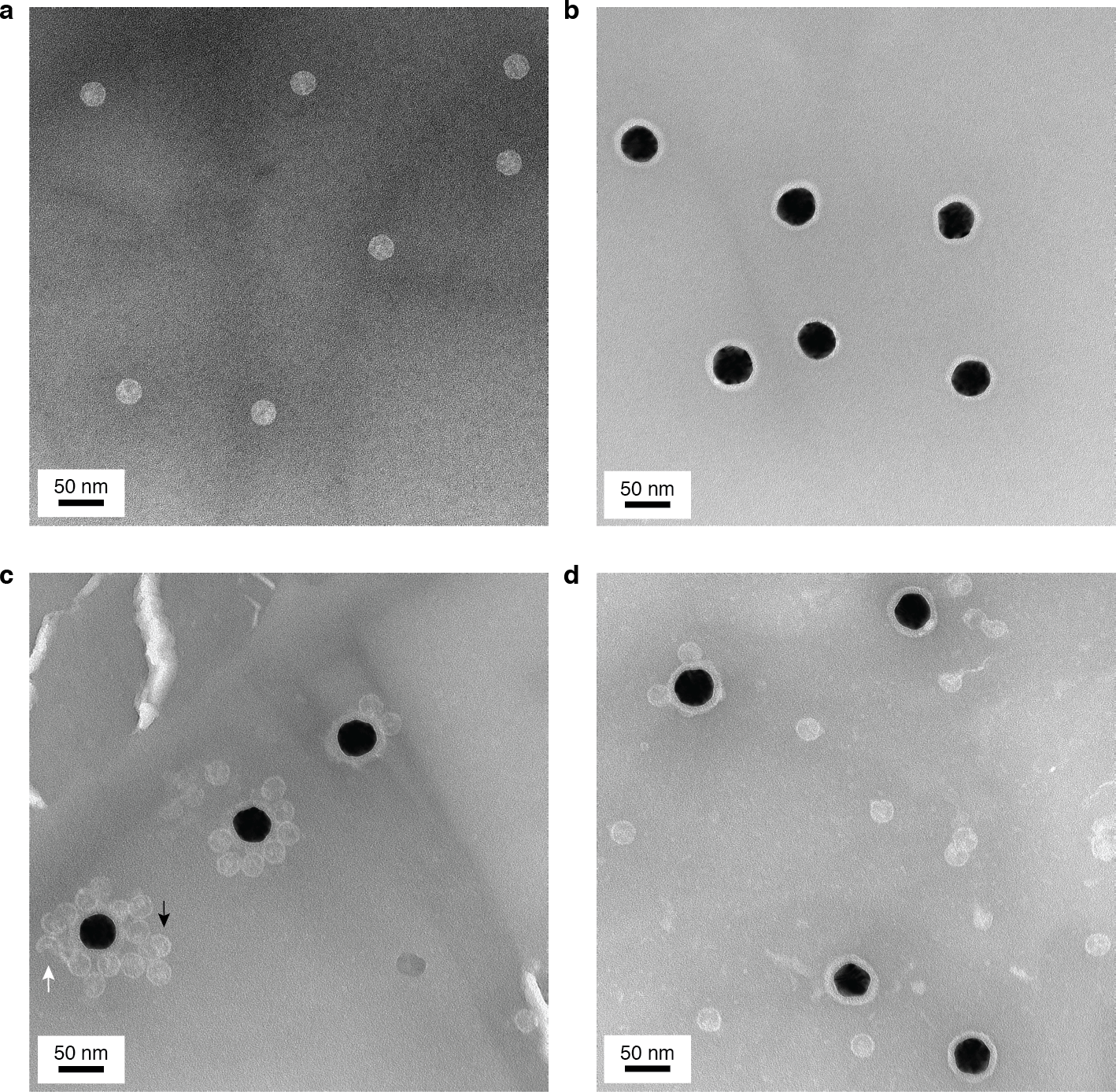
Negatively stained transmission electron micrographs of virus particles, functionalized gold nanoparticles, capsids assembled around RNA strands that are bound to the surface of the gold particles, and capsids assembled around untethered RNA. Each sample is stained with methylamine tungstate stain solution (Nanoprobes) before imaging. (a) Wild-type MS2 particles. (b) Amine-functionalized 30-nm gold nanoparticles (Nanopartz) that are coated with PEG and decorated with surface oligos. The dark spots are the gold particles, and the surrounding lighter halos are the negatively stained coatings on the particle surfaces. These coatings consist of a proprietary polymer base layer, which is applied by the manufacturer to the gold nanoparticles, and the PEG-DNA molecules that we conjugate to the particles. (c) An assembly reaction in which 2 μM coat-protein dimers in assembly buffer is added to RNA-DNA complexes that have been incubated for 1 h with the functionalized gold particles. White arrow points to a partial capsid. Black arrow points to a particle that is larger than a capsid. (d) A control reaction in which 2 μM coat-protein dimers in assembly buffer is added to bare RNA that has been incubated for 1 h with the functionalized gold particles. The higher number of capsids near the surface of the gold particles for the experiments using RNA-DNA complexes suggests that these capsids assembled around RNA-DNA complexes that were tethered to the particle surface.

**EXTENDED DATA FIGURE E5.**
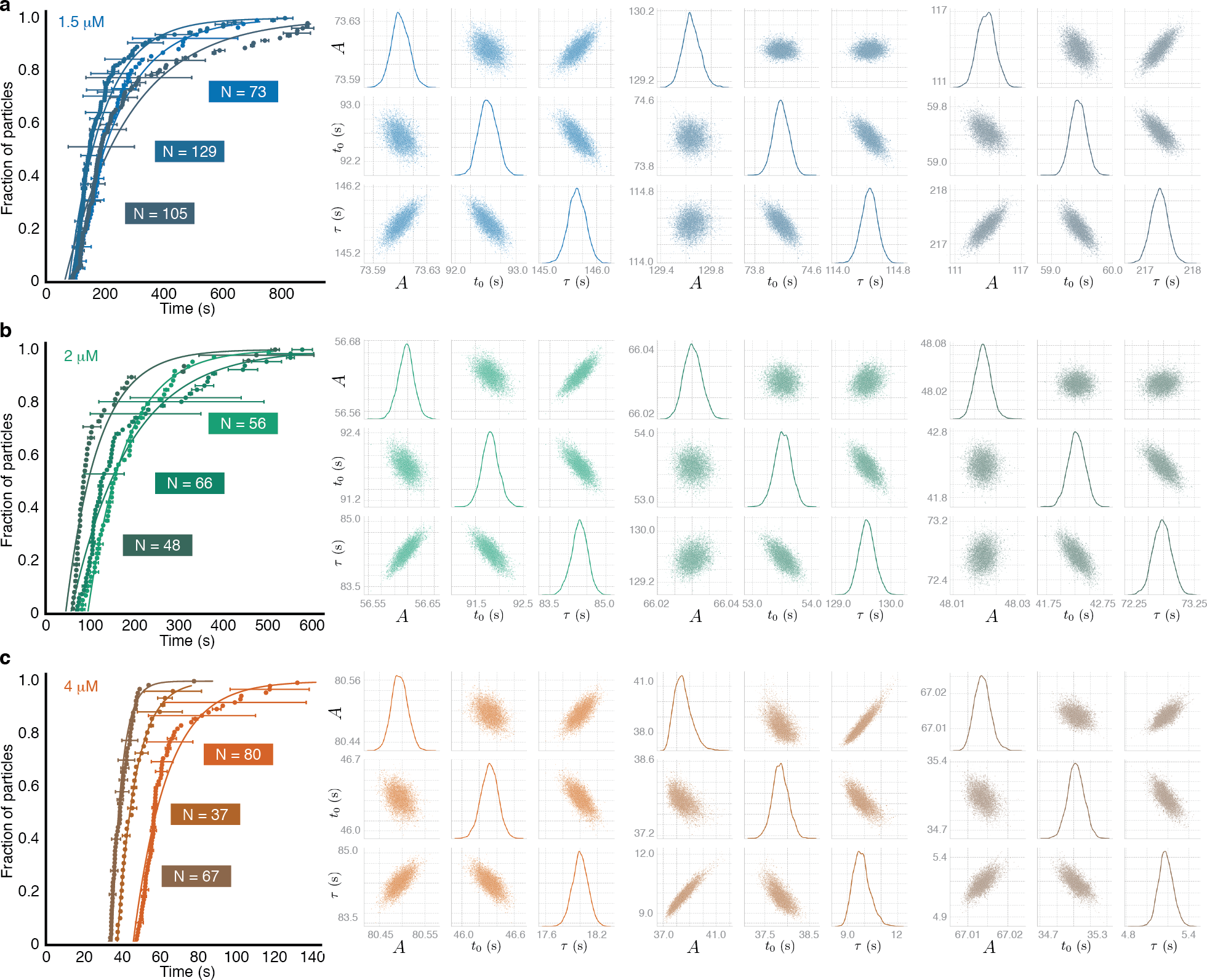
Cumulative distributions of start times and posterior probability distributions of parameter values obtained by fitting the distributions. The results from triplicate assembly experiments with (a) 1.5, (b) 2, and (c) 4 μM dimers are shown. Each cumulative distribution of start times (left) is measured from a separate assembly experiment. Uncertainties in the time measurements are represented by horizontal bars. Fits are shown as solid curves. Number of particles (N) are shown on the plot. Posterior probability distributions of parameter values (right) are sampled using a Markov-chain Monte Carlo technique. The plots along the diagonal show kernel density estimates of the fully marginalized posterior distributions of each parameter, while the off-diagonal plots show the joint distributions. The data and fit shown in the lightest color of each panel are from the experiments shown in Figs. 2, 3a, 3c, and Extended Data Figs. 7 and 8. Data from all 9 of the experiments in this figure were used to obtain the nucleation times shown in Fig. 3b.

**EXTENDED DATA FIGURE E6.**
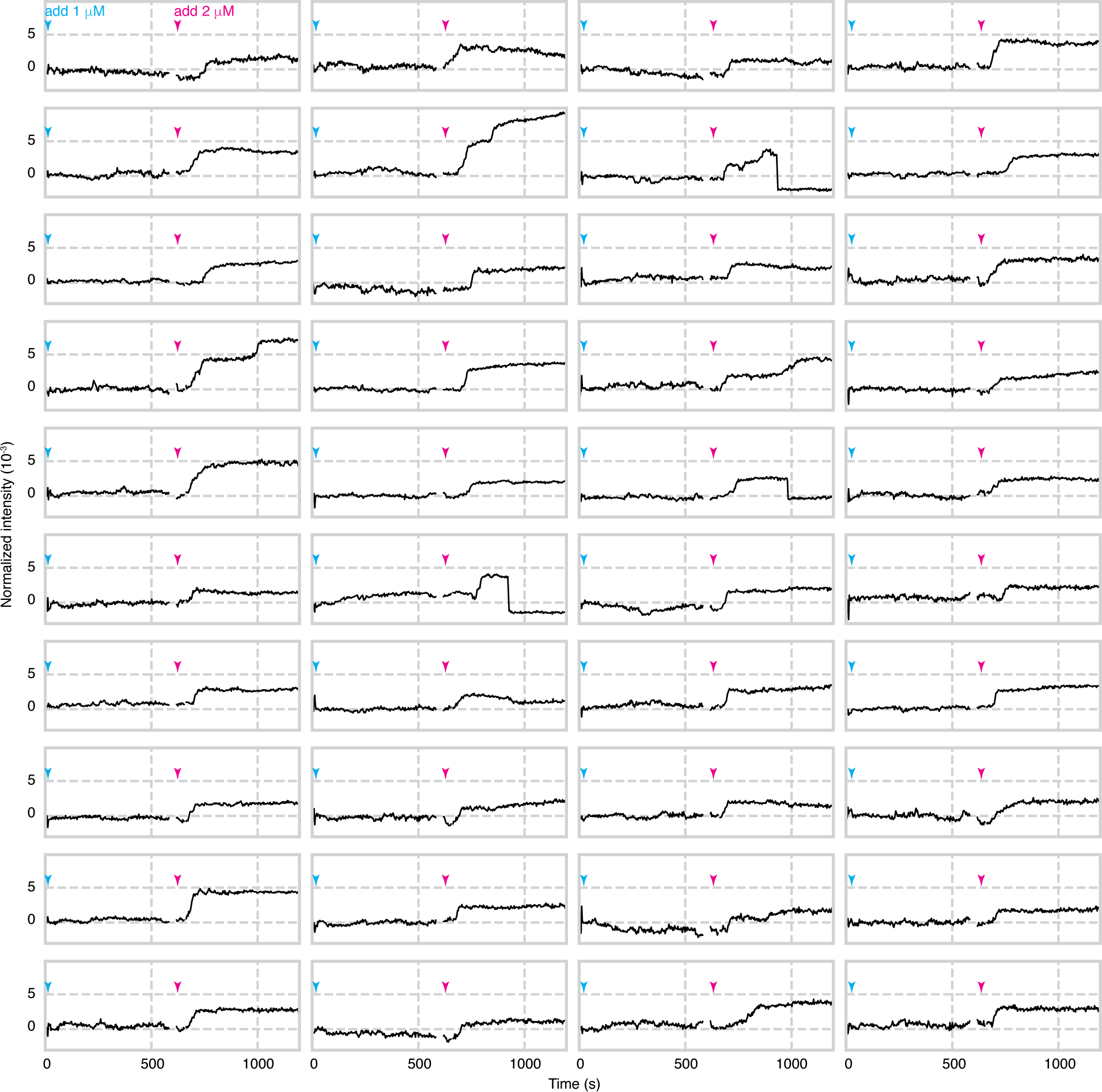
Assembly of 1 μM coat-protein dimers. When 1 μM coat-protein dimers is added (cyan arrowheads) to the surface-bound RNA, no assembling particles appear over the course of 600 s. At this point, 2 μM coat-protein dimers is added (pink arrowheads), after which we observe particles assembling at 75 locations within the field of view. Intensity traces for 40 of these particles are shown above. We also show traces for the first 600 s at the same locations. There is no data between 586 and 615 s, during which time we block the illumination beam and inject the 2 μM protein. As in Extended Data Fig. 2, we interpret abrupt drops in intensity after assembly as detachment events. The traces are measured from the data shown in Supplementary Movie 2. The data are recorded at 100 Hz and are plotted with a 300-frame average.

**EXTENDED DATA FIGURE E7.**
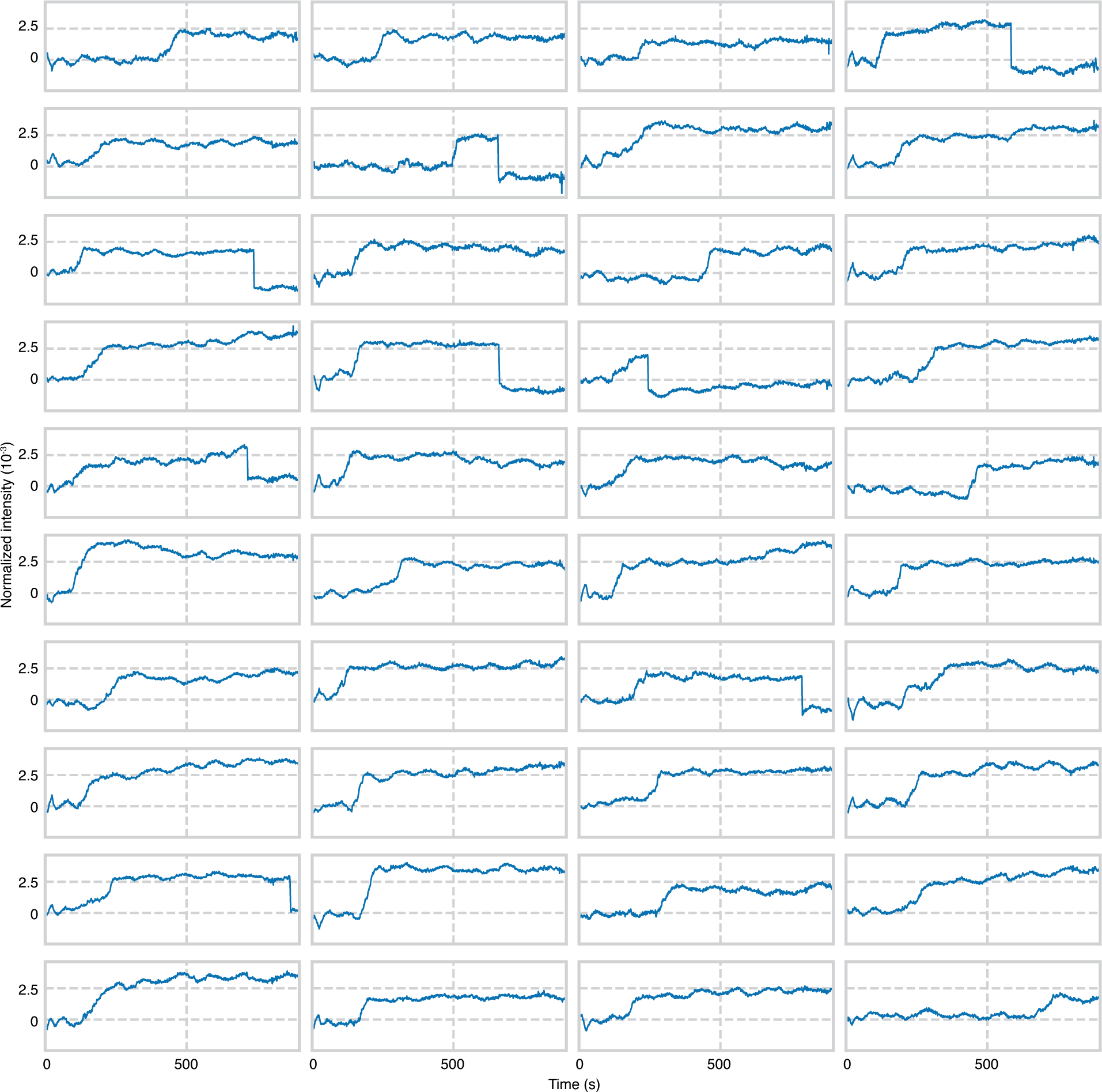
Assembly of 1.5 μM coat-protein dimers. Kinetic traces for 40 of the 73 observed assembling particles in one experiment are shown. As in Extended Data Fig. 2, we interpret abrupt drops in intensity after assembly as detachment events. A portion of the traces from the same experiment appear in Fig. 3. The final intensities of the particles in this experiment are used for Fig. 3c. The traces are measured from the data shown in Supplementary Movie 3. The data are recorded at 1,000 Hz and are plotted with a 1,000-frame average.

**EXTENDED DATA FIGURE E8.**
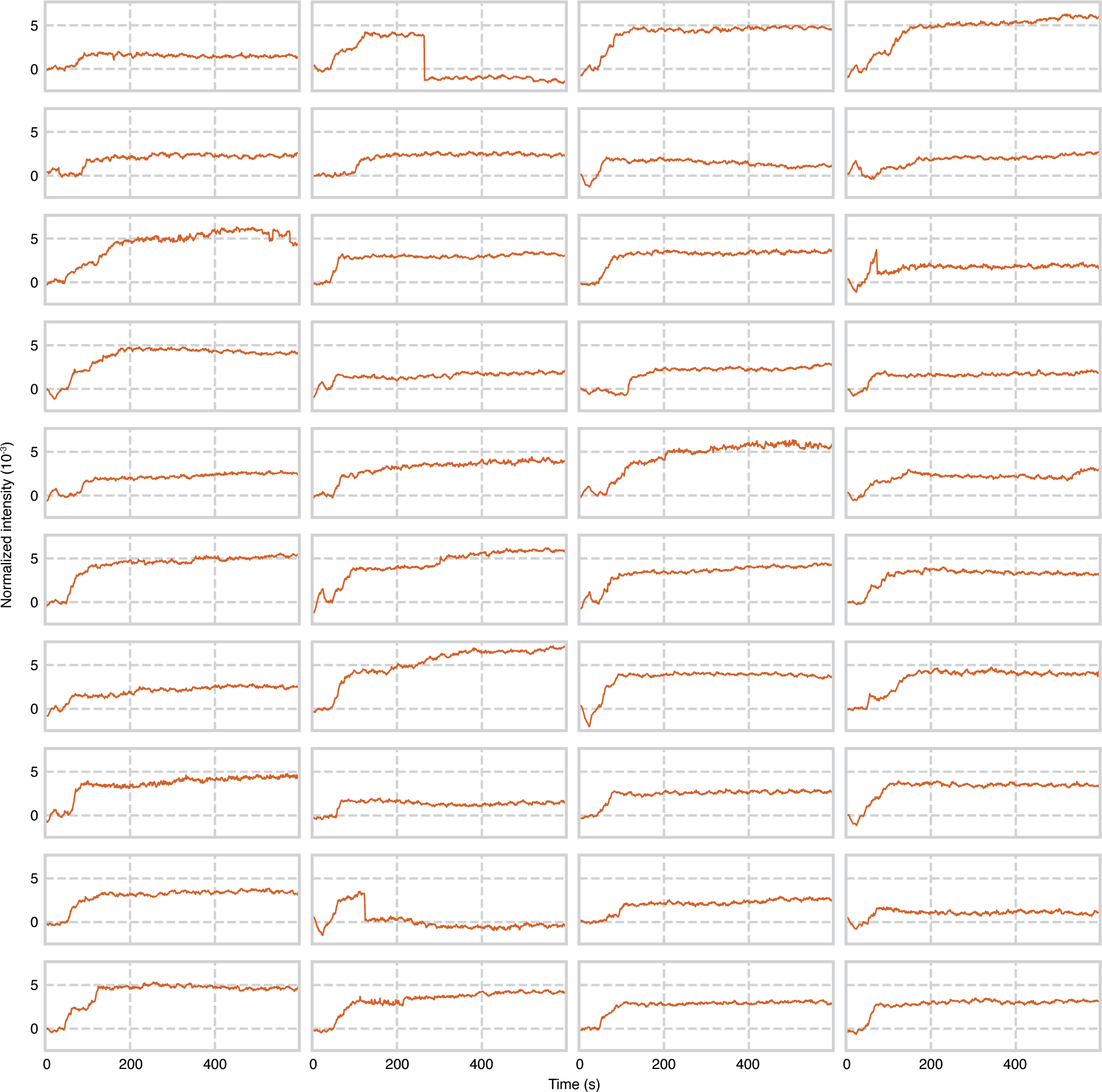
Assembly with 4 μM coat-protein dimers. Kinetic traces for 40 of the 80 observed assembling particles in one experiment are shown. As in Extended Data Fig. 2, we interpret abrupt drops in intensity after assembly as detachment events. A portion of the traces from the same experiment appear in Fig. 3. The final intensities of the particles in this experiment are used for Fig. 3c. The traces are measured from the data shown in Supplementary Movie 4. The data are recorded at 1,000 Hz and are plotted with a 1,000-frame average.

**EXTENDED DATA FIGURE E9.**
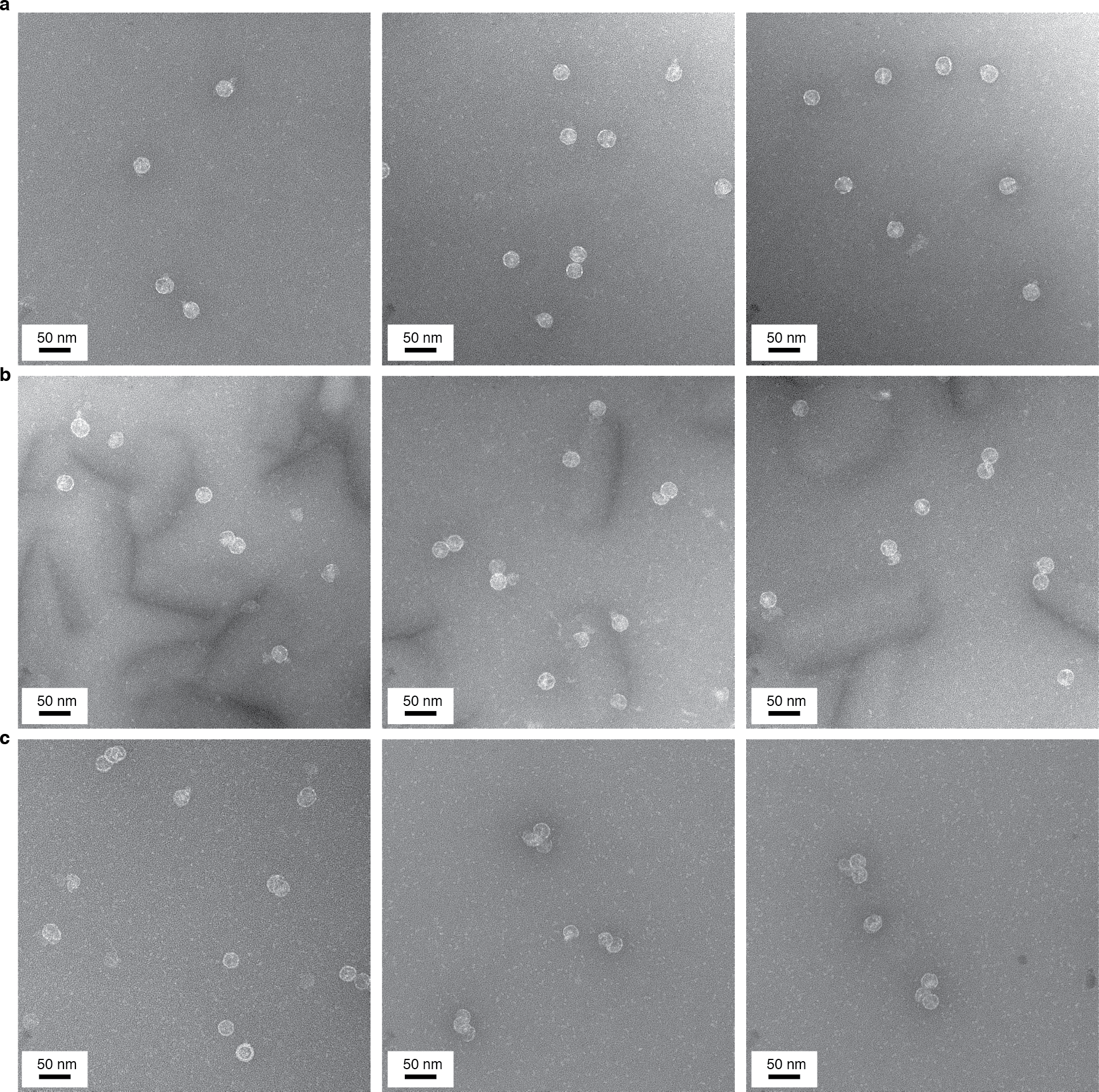
Negatively stained transmission electron micrographs of assembled particles from control experiments performed with varying concentrations of protein and untethered RNA. Each sample is stained with methylamine tungstate stain solution (Nanoprobes) before imaging. (a) Three micrographs of particles taken 20 min after mixing 1.5 μM coat-protein dimers and 10 nM RNA in assembly buffer. (b) Three micrographs of particles taken 10 min after mixing 2 μM coat-protein dimers and 10 nM RNA in assembly buffer. (c) Three micrographs of particles taken 10 min after mixing 4 μM coat-protein dimers and 10 nM RNA in assembly buffer.

